# Assessing sprint technique with shoe-mounted inertial sensors

**DOI:** 10.1101/2024.05.06.592662

**Authors:** Gerard Aristizábal Pla, Douglas N. Martini, Michael V. Potter, Wouter Hoogkamer, Stephen M. Cain

## Abstract

Negative foot speed (i.e., the speed of the backward and downward motion of the foot relative to the body at ground contact) is a strong predictor of sprinting performance. Inertial measurement units (IMUs) are becoming a popular approach for assessing sports performance. The primary aim of this study was to use IMUs to investigate the relationship between negative foot speed and top running speed attained during a sprint on an outdoor track. Seventeen participants performed 80-meter track sprints while wearing a shoe-mounted IMU. Anteroposterior and vertical components of negative foot speed were extracted from the IMU. For the mean peak stride speed of 7.98±0.78m/s, the adjusted R^2^ values were 0.27 and 0.42 versus the anteroposterior and vertical components of negative foot speed, respectively. In conclusion, our findings support the common coaching tip of increasing negative foot speed to improve sprint speed. In addition, the results of this study support the use of IMUs for quantifying sprinting technique with actionable metrics.

## Introduction

High-speed running is a crucial factor influencing performance in both individual [1] and team sports [2,3]. Thus, a significant objective in sport training is improving sprint performance. Kinetic determinants of sprint performance include the horizontal component of the ground reaction forces (GRF) or the ratio of force (i.e., the orientation of the GRF vector in the sagittal plane) [4,5,6,7]. Kinematic determinants of sprint performance include spatiotemporal parameters (e.g., top speed [1], contact time [8,9,10,11], step frequency [8,9,11]) and leg angular velocity [9].

Most traditional approaches to quantifying sprint performance (e.g., motion capture systems [9,11], resistance devices [12,13]) utilize equipment that is expensive or that have a very specific and inflexible setup, making it challenging for coaches and athletes to implement measurements of sprint biomechanics in day-to-day training. Wearable inertial measurement units (IMUs) offer a cost-effective and user-friendly alternative for assessing sprinting performance technique. These wearables are small (e.g., 42×27×11mm) and generally comprise tri-axial accelerometers and gyroscopes, that measure linear accelerations and angular rates, respectively [14].

Shoe-mounted IMUs have been used to estimate sprinting performance determinants [15,16,17]. Martín-Fuentes et al. [15] found that plantarflexion velocity showed the greatest association with sprint performance and that ground contact time was also associated with sprint performance, with faster sprinters running with shorter ground contact time. Shoe-mounted IMUs can also be utilized to estimate foot kinematics for sprinting with the zero-velocity update (ZUPT) method with good validity for peak sprint speeds of up to 8.00±0.88 m/s [16, 17]. In particular, IMUs and the ZUPT method provide accurate estimates of stride length and cumulative distance traveled for sprinting speeds [16].

The ZUPT method allows for the calculation of stride velocities, stride lengths and stride times. Stride lengths and times can be used to obtain estimates of the runner’s speed. Computation of the instantaneous speed of the center of mass (COM) for every stride of a sprint results in a velocity curve [16]. Then, the method proposed by Samozino et al. [18] can be used with split times as inputs to obtain force and power outputs (e.g., ratio of force, horizontal power). ZUPT implementation for fast running requires the detection of stance phases [16, 19], which enables the calculation of various spatiotemporal metrics (e.g., contact time, swing time, step frequency). IMUs can therefore be used to assess sprint performance [16]. However, these data are less actionable for coaches and sprinters. For example, faster speeds are correlated with shorter ground contact times [8,10,9,11], but this information alone may not provide specific actionable steps or interventions to improve an athlete’s sprint performance. One common actionable approach by coaches for improving sprint performance focuses on increasing negative foot speed (i.e., the speed of the backward and downward motion of the foot relative to the body at ground contact) [11,20,21,22].

The increase in negative foot speed referred as the “pawing” or “shipping” action of the foot, is commonly coached in athletics [20]. Several studies [11,21,22] have found significant correlations between the anteroposterior (AP) component of negative foot speed (i.e., the relative velocity vector at touchdown: the velocity vector at touchdown with respect to the runner’s speed), and peak running speed. Haugen et al. [11] found significant correlations between the AP component of negative foot speed and peak running speed in an indoor track for 20-meter flying sprints using a motion capture system. Murphy et al. [22] found significant correlations between the AP component of negative foot speed and peak running speed in an outdoor track for 40-60-meter sprints using high-speed cameras. Clark et al. [21] found the strongest correlations between the AP component of negative foot speed and average peak running speed during the last 8 meters of a 40-meter sprint in an indoor athletic facility using a motion capture system. Therefore, the AP component of negative foot speed, measured with different devices and in different running environments, is a strong predictor of sprinting performance [11,21,22]. With IMUs becoming more popular in sports performance applications, we set out to evaluate the relationship between negative foot speed and top-speed sprinting.

The main aim of this study was to use IMUs to investigate the relationship between the IMU-calculated AP component of negative foot speed and peak sprinting speed attained during an 80-meter sprint on an outdoor track. Based on optical motion capture or high-speed camera studies [11,21,22], we hypothesized that faster peak speeds would be achieved with greater AP components of negative foot speed. We additionally investigated the relationship between the vertical (VT) component of negative foot speed and peak sprinting speed. Only one study [22] examined the relationship between the VT component of negative foot speed and peak sprinting speed but did not find significant correlations. Therefore, more studies examining the VT component of negative foot speed are needed to better assess the VT component of negative foot speed as being a predictor of sprinting performance.

## Methods

### Participants

This study involved seventeen participants (13 males, 4 females), each over the age of 18 years; recruitment period: March 14, 2022 – May 25, 2022. Summary descriptive characteristics for all participants are listed in Table 1 [16]. Inclusion criteria were the ability to run at a speed of 7m/s or faster and no injuries or surgeries for at least three months prior to the testing session. Exclusion criteria were orthopedic, cardiovascular, or neuromuscular conditions that could potentially affect sprint performance. All participants provided written informed consent that was approved by the Institutional Review Board of the University of Massachusetts Amherst (IRB protocol number 3143).

**Table 1.**
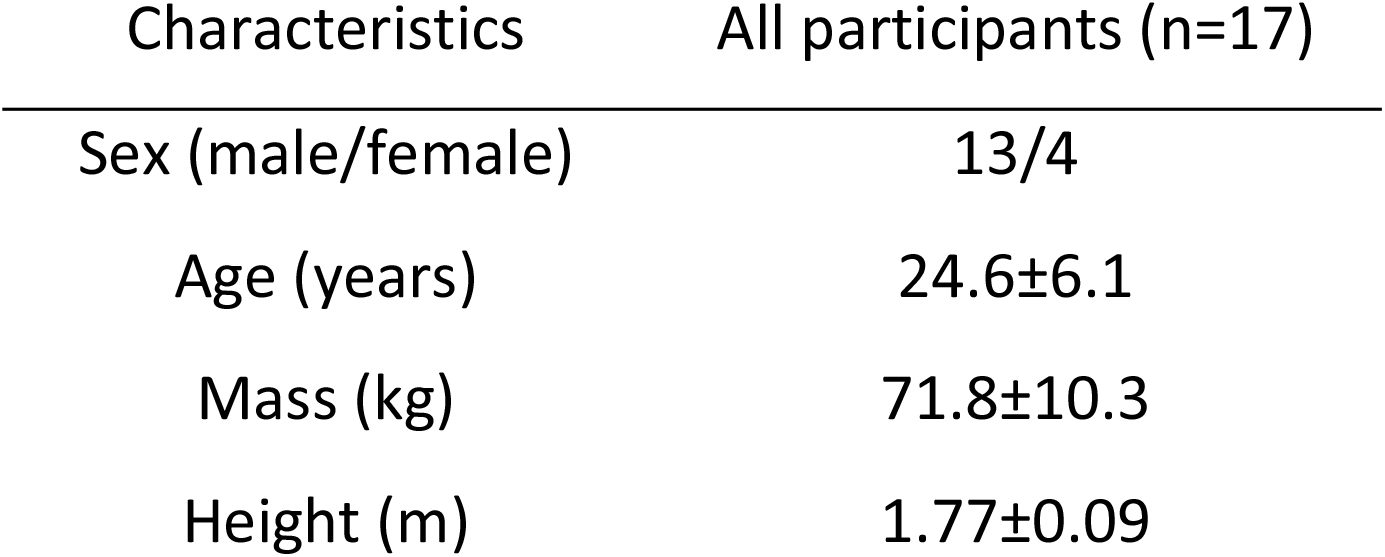
Descriptive characteristics for all participants.

| Characteristics | All participants (n=17) |
| --- | --- |
| Sex (male/female) | 13/4 |
| Age (years) | 24.6±6.1 |
| Mass (kg) | 71.8±10.3 |
| Height (m) | 1.77±0.09 |

### Experimental protocol

The experimental protocol has been explained previously [16]. Briefly, an IMU (low-g ±16 g range, high-g ± 200 g range, ± 2000 deg/s range, sampling at 1125Hz, 16-bit resolution, mass=9.5g; Blue Trident ImeasureU, Vicon Motion Systems Ltd, Oxford, UK) was affixed to the right shoe’s medial dorsal area [19] using double-sided tape and secured with Hypafix (BSN Medical, Hamburg, Germany) tape to minimize motion artefacts (Fig 1). After a self-selected warm-up, each participant performed an 80-meter sprint at maximum effort at an outdoor track. Participants were instructed to maintain a stationary position for approximately 15 seconds prior to commencing the sprint, with the IMU aligned directly over the start line.

**Fig 1.**
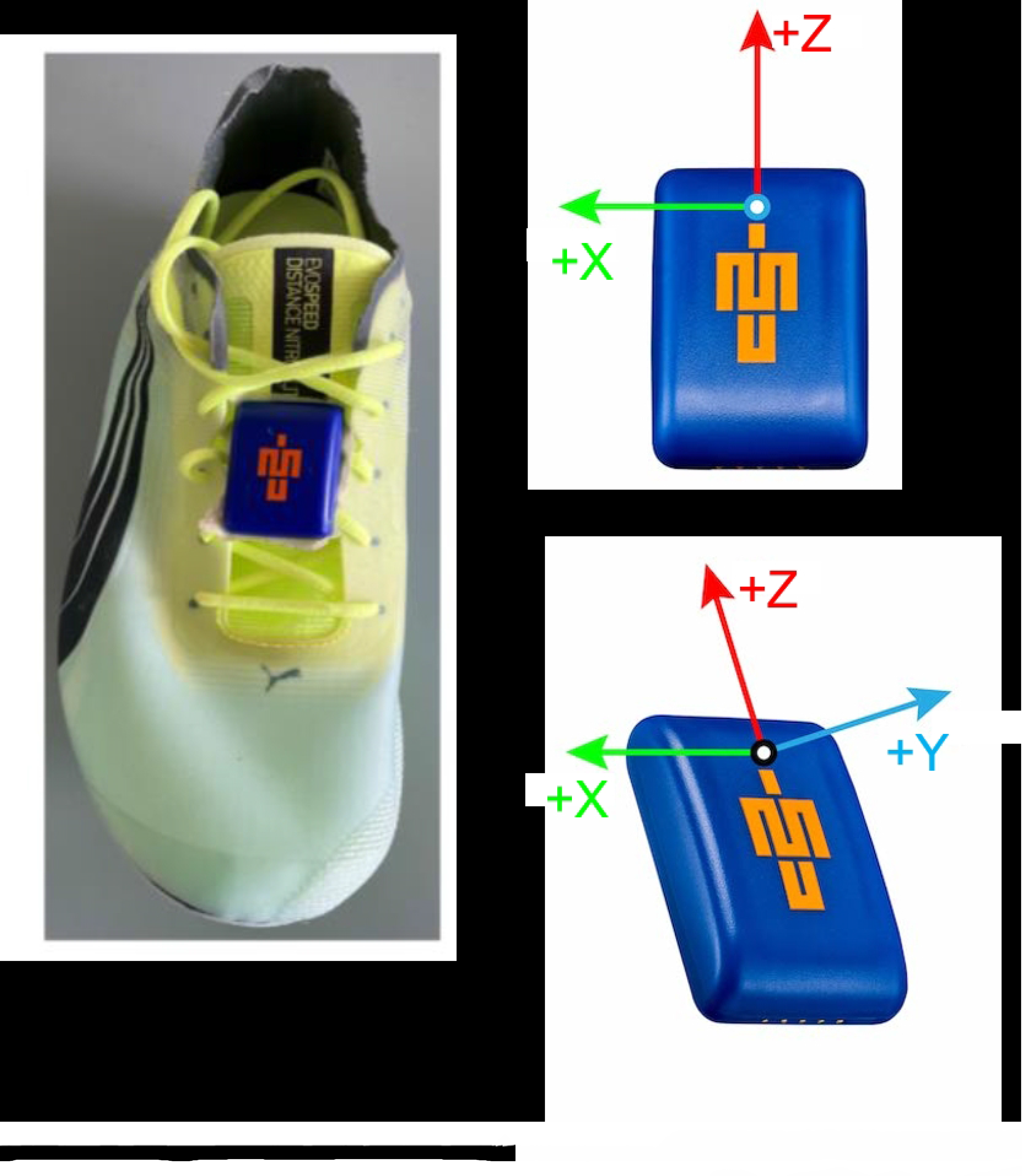
Blue Trident IMeasureU attachment to the instep of the shoe.

### IMU analysis

We analyzed the raw IMU data using customized software in Python (Python Software Foundation, Delaware, USA). To address saturation in the low-g accelerometer, the low-g and high-g accelerometers were synchronized using down sampling and cross correlation analysis. The low-g accelerometer signal was used when it was not saturated (linear accelerations smaller than ±16g) and the high-g accelerometer signal was used when the low-g accelerometer saturated [16].

The IMU-fixed frame measurements were rotated to a foot-fixed frame with axes aligned with anatomical directions using a procedure similar to [23]. The average IMU-measured three-dimensional acceleration due to gravity measured during the ∼15 seconds of standing still 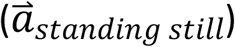 was used to define a foot vertical axis 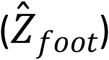 aligned with gravity:

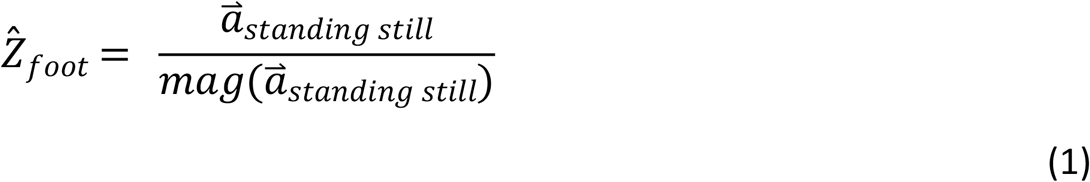

where *mag* denotes the vector magnitude. A foot mediolateral axis 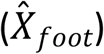 was defined as an orthogonal unit vector to 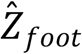 and the IMU orthogonal unit negative Z vector ([0,0, ― 1]) (Fig 1):

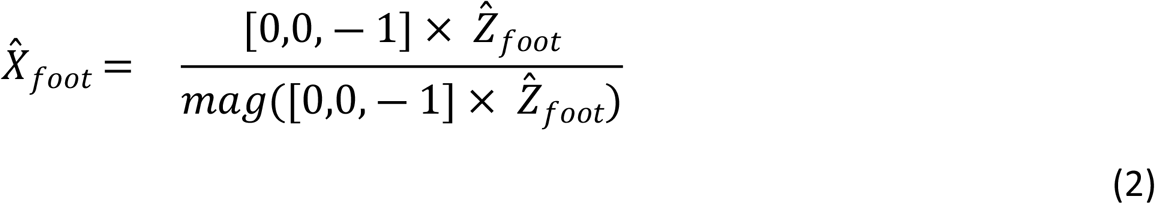

where × denotes the cross product. A foot anteroposterior axis (*Ŷ*_*food*_) was defined as an orthogonal unit vector to 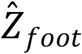 and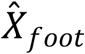:

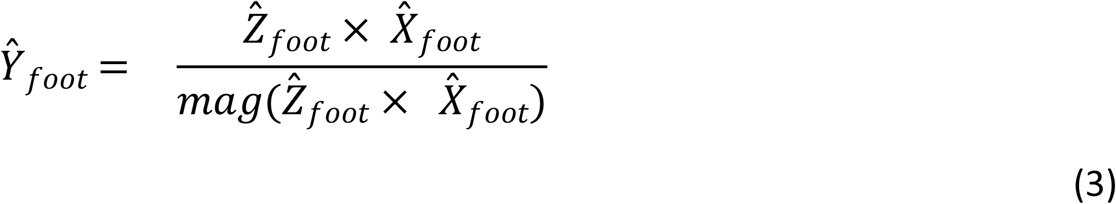

The resulting orthogonal vectors 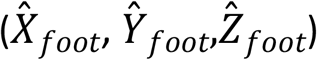 were then used to define a rotation matrix that rotates the IMU measurements from the IMU frame to the foot frame.

Next, the ZUPT method was implemented and stride velocities, stride lengths and stride times were obtained [16]. Time points at foot contact, defined when the VT acceleration signal with gravity subtracted changed from negative to positive prior to maximal peaks in the acceleration magnitude, were identified (Fig 2).

**Fig 2.**
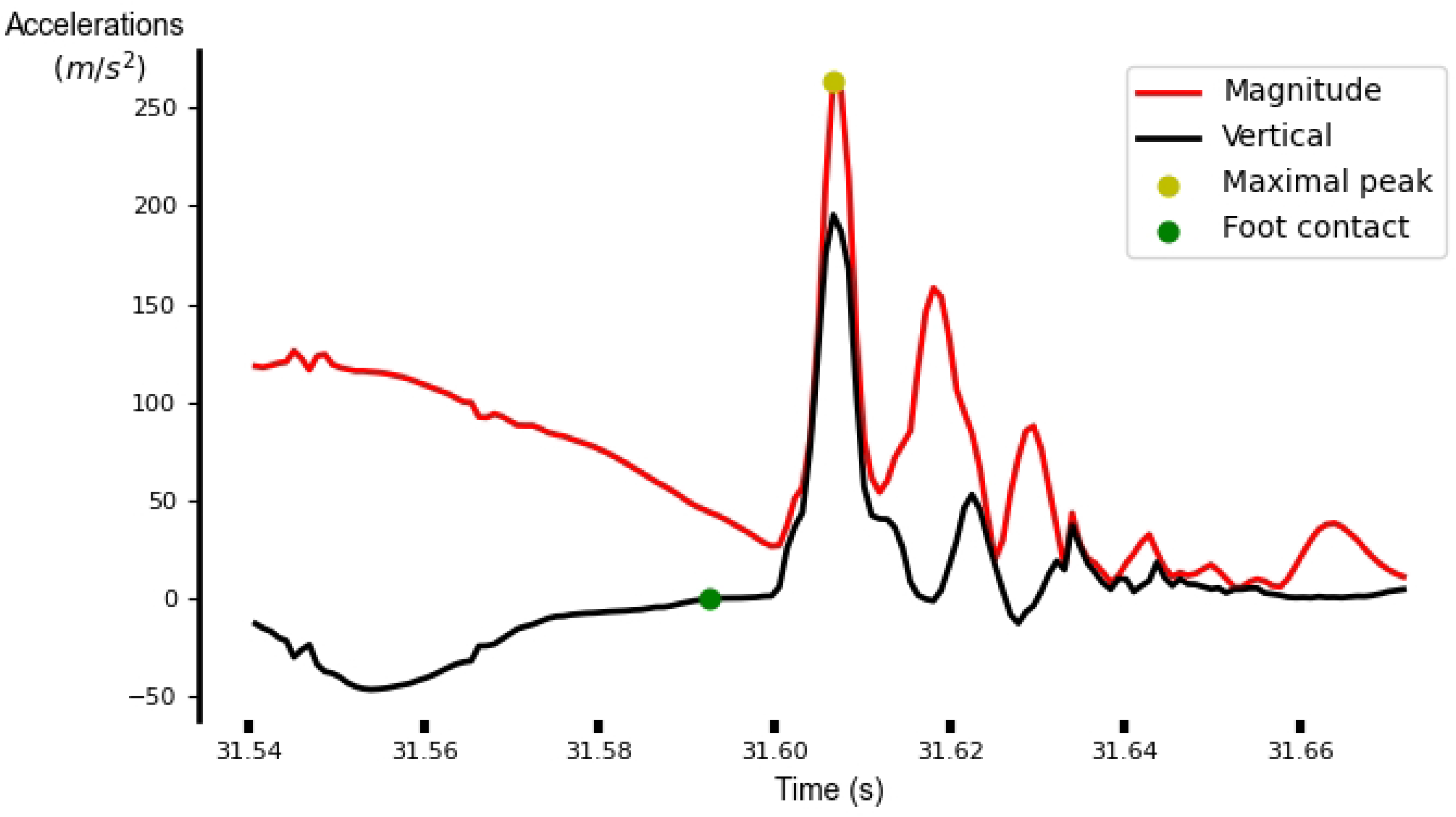
Sample identification of time points at touchdown.

Since stride velocities are affected by yaw drift, a yaw rotation was applied to align stride velocities with the direction of running. Following this rotation, for a given stride, the AP-axis of the stride reference frame was defined by the displacement of the foot during the stride, the VT-axis was aligned with vertical and the mediolateral axis was formed by the cross product of VT and AP.

Estimates of runner’s stride speeds were obtained by dividing stride lengths by stride times. Finally, these speeds were subtracted from aligned foot velocities at the time points of touchdown to obtain the velocity vectors at touchdown. Peak sprinting speed was calculated for all sprints as the fastest stride speed. The velocity vector at the top speed stride was also computed. Therefore, for each participant three values were used for the statistical analysis (i.e., top stride speed, global VT velocity vector for the top speed stride, relative AP velocity vector for the top speed stride).

### Statistical analysis

Rstudio (R Core team, Auckland, New Zealand) was used for the statistical analysis. We used simple linear regressions to investigate the relationship between the velocity vector and top speed. Top stride speeds were included as dependent variables and the VT and AP components of the velocity vector as independent variables. Alpha level was set a priori to 0.05 for the slope of the regression and the confidence intervals.

## Results

### Effect of sprinting speeds and body mass on stride length estimation

The relative AP velocity vector had a significant relationship to peak stride speed. Faster peak speeds were achieved with greater AP components of the relative velocity vector (R^2^=0.27; p=0.02; y=-0.8-0.54x [where x denotes top stride speed]; Fig 3; Table 2). The global VT velocity vector had a significant relationship to peak stride speed. Faster peak speeds were achieved with greater VT components of the global velocity vector (R^2^=0.42; p=0.003; y=0.2-0.37x [where x denotes top stride speed]; Fig 4; Table 2).

**Table 2.** Simple linear regression results. Significant differences are highlighted in bold.

| Velocity vector component | *p* value | Adjusted R<sup>2</sup> value | Standardized $\beta$ speed | Confidence intervals | Linear equation [x denotes top stride speed] |
| --- | --- | --- | --- | --- | --- |
| Anteroposterior | **0.018** | 0.27 | -0.57 | (-0.97, -0.11) | $y = -0.8 - 0.54x$ |
| Vertical | **0.003** | 0.42 | -0.68 | (-0.59, -0.15) | $y = 0.2 - 0.37x$ |

**Fig 3.**
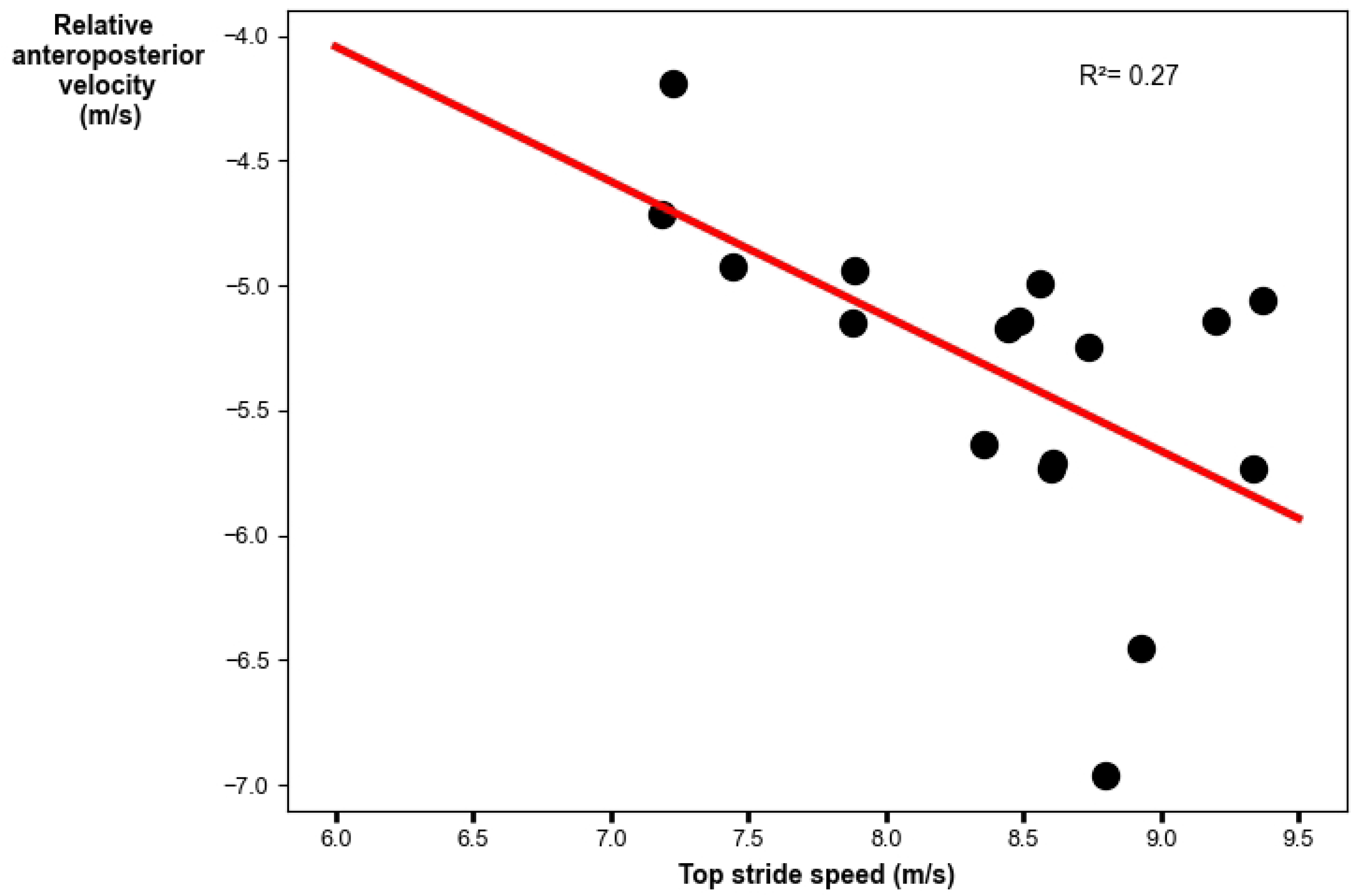
Linear regression analysis for the relative anteroposterior velocity vector with peak stride speed. Faster top speeds were achieved with greater magnitude (larger negative) anteroposterior components of the relative velocity vector.

**Fig 4.**
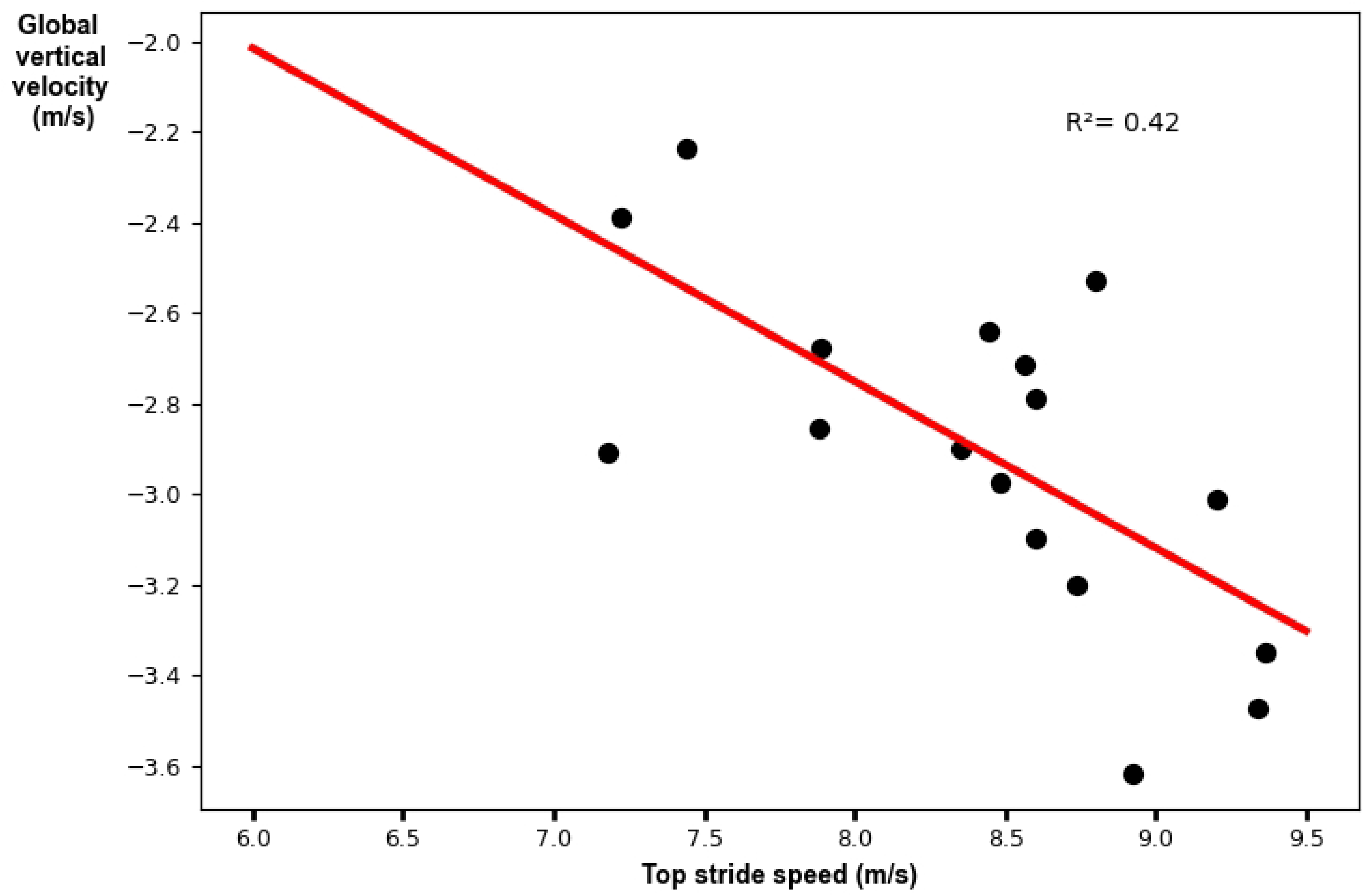
Linear regression analysis for the global vertical velocity vector with peak stride speed. Faster top speeds were achieved with greater magnitude (larger negative) vertical components of the global velocity vector.

## Discussion

The main aim of this study was to investigate the relationship between negative foot speed and peak running speed attained during an 80-meter sprint. We found that the AP and VT components of negative foot speed were significantly correlated to peak stride speed. Specifically, for peak stride speeds of 7.98±0.78m/s the adjusted R^2^ values were 0.27 and 0.42 for the AP and VT component of negative foot speed, respectively. Our findings align with prior studies, confirming that the AP component of negative foot speed is an important determinant of sprinting performance [11,21,22]. Our findings also demonstrate that the VT component of negative foot speed is an important determinant of sprinting performance. This kinematic parameter, including both components, is of interest to coaches and its increase is referred as the “pawing” or “shipping” action of the foot [20]. Thus, our findings support the common coaching tip of the “pawing” or “shipping” action of the foot to improve sprint performance.

Our findings also build further on limited available research for assessing sprinting technique with actionable metrics using only shoe-mounted IMUs [15]. Martín-Fuentes et al. [15] found that plantarflexion velocity correlated most strongly with sprint performance and shorter ground contact times were associated with faster sprinters. We found that faster peak speeds were achieved with greater VT and AP components of negative foot speed. Therefore, we recommend the use of shoe-mounted IMUs for sprint technique evaluation. In addition, shoe-mounted IMUs have several advantages over traditional approaches to quantify sprint performance technique, such as cost-effectiveness, ease of set-up, minimal interference with the running movement, and the ability to deliver instant feedback. Therefore, our work may further increase the utility of IMUs for sprint coaching and performance evaluation over traditional approaches to quantify sprint technique.

Certain limitations must be acknowledged in the present study. First, IMU-based negative foot speed was not compared against the ground truth negative foot speed measured by an optical motion capture system. While previous work has validated other kinematic quantities obtained from shoe-mounted IMUs during sprinting (e.g., stride length [16, 17], contact time [24, 25]), future work should assess the validity of IMU-based estimates of instantaneous foot speed against optical motion capture. Second, IMU-based peak speeds have more uncertainty than speeds measured by an optical motion capture system [16]. Thus, while the use of IMUs to measure foot speed is more practical for implementation in coaching scenarios, future work should aim to confirm the relationships between negative foot speed and peak speed observed in this work using gold-standard optical motion capture.

In conclusion, the results of this study support the use of IMUs for quantifying sprinting technique with actionable metrics. Faster peak speeds were achieved with greater VT and AP components of negative foot speed at touchdown.

## Acknowledgments

The authors express gratitude to the members of the UMILL lab who assisted in data collection. Special thank you to Dr. Alex Shorter and Dr. Loubna Baroudi for sharing their insights, and Dr. Leia Stirling for her assistance in software development and ZUPT implementation.

